# PEPc-mediated CO_2_ assimilation provides carbons to gluconeogenesis and the TCA cycle in both dark-exposed and illuminated guard cells

**DOI:** 10.1101/2021.11.03.467183

**Authors:** Valéria F. Lima, David B. Medeiros, Silvio A. Cândido-Sobrinho, Francisco Bruno S. Freire, Nicole P. Porto, Alexander Erban, Joachim Kopka, Markus Schwarzländer, Alisdair R. Fernie, Danilo M. Daloso

## Abstract

Evidence suggests that guard cells have higher rate of phospho*enol*pyruvate carboxylase (PEPc)-mediated dark CO_2_ assimilation than mesophyll cells. However, it is unknown which metabolic pathways are activated following dark CO_2_ assimilation in guard cells. Furthermore, it remains unclear how the metabolic fluxes throughout the tricarboxylic acid (TCA) cycle and associated pathways are regulated in illuminated guard cells. Here we used ^13^C-HCO_3_ labelling of tobacco guard cells harvested under continuous dark or during the dark-to-light transition to elucidate principles of metabolic dynamics downstream of CO_2_ assimilation. Most metabolic changes were similar between dark-exposed and illuminated guard cells. However, illumination increased the ^13^C-enrichment in sugars and metabolites associated to the TCA cycle. Sucrose was labelled in the dark, but light exposure increased the ^13^C-labelling into this metabolite. Fumarate was strongly labelled under both dark and light conditions, while illumination increased the ^13^C-enrichment in pyruvate, succinate and glutamate. Only one ^13^C was incorporated into malate and citrate in either dark or light conditions. Our results collectively suggest that the PEPc-mediated CO_2_ assimilation provides carbons for gluconeogenesis, the TCA cycle and glutamate synthesis and that previously stored malate and citrate are used to underpin the specific metabolic requirements of illuminated guard cells.

**Highlight:** PEPc-mediated CO_2_ assimilation provides carbons for gluconeogenesis and the TCA cycle, whilst previously stored malate and citrate are used to underpin the specific metabolic requirements of illuminated guard cells.

## Introduction

Guard cell metabolism has been studied for over a century. From the starch-sugar conversion hypothesis to the discovery that inorganic ions such as potassium (K^+^) modulate guard cell osmotic potential during light-induced stomatal opening (Lloyd, 1908; Fischer, 1968), the understanding of how guard cell metabolism regulates stomatal movements has evolved dramatically over the decades (Granot and Kelly, 2019). Breakdown of sugars, starch and lipids within guard cells are important mechanisms to sustain the speed of light-induced stomatal opening (Antunes *et al*., 2012; Ni, 2012; Daloso *et al*., 2015*a*, 2016*a*; Horrer *et al*., 2016; McLachlan *et al*., 2016; Medeiros *et al*., 2018*a*; Flütsch *et al*., 2020*a*). Given the sink characteristics of guard cells (Hite *et al*., 1993; Ritte *et al*., 1999), the import of ATP, organic acids and sugars from mesophyll cells contributes to the stomatal opening upon illumination (Hedrich and Marten, 1993; Araújo *et al*., 2011*a*; Kelly *et al*., 2013; Wang *et al*., 2014*a*, 2019; Lugassi *et al*., 2015; Medeiros *et al*., 2016; Antunes *et al*., 2017; Flütsch *et al*., 2020*b*). Thus, light mediates stomatal opening by both, perception directly in the guard cells (Kinoshita *et al*., 2001; Wang *et al*., 2010; Ando and Kinoshita, 2018) and indirectly through signals derived from mesophyll cells (Mott, 2009; Fujita *et al*., 2019; Flütsch and Santelia, 2021). For the latter mode the respiratory activity seems to be of great importance (Nunes-Nesi *et al*., 2007*a*; Araújo *et al*., 2011*a*; Vialet□Chabrand *et al*., 2021). Taken together, these works have substantially improved our understanding of light-induced changes in guard cell metabolism. However, light is just one out of several stimuli that affects stomatal aperture and how guard cell metabolism operates in the dark remains unexplored.

As the main energy source for plants, folial illumination activates photosynthesis alongside other mechanisms that sustain plant metabolism (Buchanan, 2016). Photosynthetic ATP production and CO_2_ assimilation mediated by ribulose-1,5-bisphosphate carboxylase/oxygenase (RuBisCO) are pivotal for the functioning of photoautotrophic cells. This is especially true for C3 and C4 plants, in which the vast majority of carbon assimilation occurs in the light period (Hohmann-Marriott and Blankenship, 2011; Matthews *et al*., 2020). Additionally, given the capacity of plant cells to store and transport photoassimilates, the remobilisation of photosynthesis-derived products such as starch and sucrose is an important mechanism that sustains both metabolism and growth during the night period (Sulpice *et al*., 2014; Apelt *et al*., 2017; Mengin *et al*., 2017). By contrast to C3- and C4-plants, a special group of plants that evolved Crassulaceae Acid Metabolism (CAM) can also fix CO_2_ through the activity of phospho*enol*pyruvate carboxylase (PEPc) in the dark (Ranson and Thomas, 1960; O’Leary *et al*., 2011). In these plants, the stomata open in the dark and close in the light. Whilst the stomatal opening in the dark favours the PEPc-mediated CO_2_ fixation, stomatal closure in the light avoids water loss, leading to the highest water use efficiency (WUE) observed in plants. Curiously, several C3 and C4 plants maintain a substantial transpiration stream during the night period (Caird *et al*., 2007; Costa *et al*., 2015). Furthermore, nocturnal stomatal conductance (*g*_sn_ - the rate of stomatal opening in the night) was positively correlated with relative growth rate in a multi-species meta-analysis (Resco de Dios *et al*., 2019). However, the regulation of *g*_sn_ remains insufficiently understood (Gago *et al*., 2020). Given the role of guard cell metabolism in regulating stomatal opening (Daloso *et al*., 2017; Lawson and Matthews, 2020), unveiling how guard cell metabolism operates in the dark will be a prerequisite for a mechanistic understanding of *g*_sn_ regulation.

Radiotracer experiments suggest that guard cells can incorporate CO_2_ in the dark at higher rates than mesophyll cells (Gotow *et al*., 1988). Furthermore, ^13^C-labelling results using mesophyll and guard cell protoplasts highlight that guard cells have faster and higher ^13^C-incorporation into malate under illuminated conditions (Robaina-Estévez *et al*., 2017). These results suggest a high PEPc activity in guard cells under either dark or light conditions. Indeed, recent ^13^C-positional labelling analysis confirmed that guard cells are able to assimilate CO_2_ in the dark, as evidenced by the increases in the ^13^C-enrichment in the carbon 4 (4-C) of malate (Lima *et al*., 2021), which is derived from PEPc activity (Abadie and Tcherkez, 2019). High dark CO_2_ fixation rates are typically found in CAM cell types (Cockburn, 1983). However, the idea that guard cells have CAM-like metabolism is not fully supported by transcriptome studies (Wang *et al*., 2011; Bates *et al*., 2012; Bauer *et al*., 2013; Aubry *et al*., 2016). It remains therefore unclear whether the metabolic photosynthetic mode of guard cells most closely resembles that of C3, C4 or CAM cells. Alternatively, guard cells might not strictly fit into any of these classifications. If this is the case, a specific mode of regulation may be predicted in these cells. This idea is supported by the finding that illumination triggers specific responses observed in guard cells, but not in mesophyll cells, such as the degradation of lipids, starch and sugars and the activation of glycolysis (Hedrich *et al*., 1985; Zhao and Assmann, 2011; Daloso *et al*., 2015*a*, 2016*a*; Horrer *et al*., 2016; McLachlan *et al*., 2016; Robaina-Estévez *et al*., 2017; Medeiros *et al*., 2018*a*; Flütsch *et al*., 2020*a*). Additionally, the metabolic fluxes throughout the TCA cycle and associated metabolic pathways seems to be differentially regulated in illuminated guard cells, when compared to mesophyll cells (Daloso *et al*., 2017; Robaina-Estévez *et al*., 2017). These observations raise the question how light influences the regulation of key metabolic pathways in guard cells.

Here we addressed this question using ^13^C-HCO_3_ labelling in guard cells subjected to either dark or light conditions. To establish cell type-specific metabolic regulation and obtain an advanced picture of the metabolic flux landscape of guard cells in the light we integrate our current data with those of illuminated Arabidopsis (*Arabidopsis thaliana* L.) rosettes following provision of ^13^CO_2_ and with illuminated Arabidopsis guard cells following provision of ^13^C-sucrose (Szecowka *et al*., 2013; Medeiros *et al*., 2018*a*).

## Material and methods

### Plant material and growth condition

Seeds of *Nicotiana tabacum* (L.) were germinated and cultivated in a substrate composed by a mixture of vermiculite, sand and soil (2:1:0.5) for 30 days. Plants were kept well-watered and nourished with Hoagland nutrient solution every week (Hoagland and Arnon, 1950) under non-controlled greenhouse conditions with natural 12 h of photoperiod, ambient temperature 30 ± 4 °C, relative humidity 62 ± 10% and photosynthetic photon flux density (PPFD) which reached a maximum value of 500 µmol photons m^-2^ s^-1^.

### Guard cell isolation

A pool of guard cell-enriched epidermal fragments (simply referred here as guard cells) was isolated following a protocol that was optimized for metabolite profiling analysis (Daloso *et al*., 2015*a*). Guard cells were isolated at pre-dawn by blending approximately three expanded leaves per replicate in a Warring blender (Philips, RI 2044 B.V. International Philips, Amsterdam, The Netherlands), that contains an internal filter to remove excess mesophyll cells, fibres and other cellular debris. The viability of the guard cells was analysed by staining with fluorescein diacetate and propidium iodide dyes, as described earlier (Huang *et al*., 1986). This analysis demonstrated that only the guard cells are alive in the epidermal fragments (Daloso *et al*., 2015*a*; Antunes *et al*., 2017). All guard cell isolations were carried out in the dark in order to maintain closed stomata and simulate opening upon illumination, following the natural stomatal circadian rhythm (Daloso *et al*., 2016*a*; Antunes *et al*., 2017).

### ^13^C-isotope labelling experiment

After isolation, guard cells were transferred to light or dark conditions and incubated in a solution containing 50 μM CaCl_2_ and 5 mM MES-Tris, pH 6.15 in the presence of 5 mM ^13^C-NaHCO_3_. The experiment was initiated at the beginning of the light period of the day. Guard cell samples were rapidly harvested on a nylon membrane (220 µm) and snap-frozen in liquid nitrogen after 0, 10, 20 and 60 minutes under light or dark conditions.

### Metabolomics analysis

Approximately 30 mg of guard cells were disrupted and transformed into a powder by maceration using mortar, pestle and liquid nitrogen. The powder was then used for metabolite extraction. The extraction and derivatization of polar metabolites were carried out as described previously (Lisec *et al*., 2006), except that 1 ml of the polar (upper) phase, instead of 150 µl, was collected and reduced to dryness. Metabolites were derivatized by methoxyamination, with subsequent addition of tri-methyl-silyl (TMS) and finally analysed by gas chromatography electron impact – time of flight – mass spectrometry (GC-EI-TOF-MS), as described previously (Lisec *et al*., 2006). The mass spectral analysis were performed using the software Xcalibur® 2.1 (Thermo Fisher Scientific, Waltham, MA, USA), as described earlier (Lima *et al*., 2018). The metabolites were identified using the Golm Metabolome Database (http://gmd.mpimp-golm.mpg.de/) (Kopka *et al*., 2005). The metabolite content is expressed as relative to ribitol (added to each biological replicate during the extraction) on a fresh weight (FW) basis. The data is reported following recommendations for metabolomics analysis (Supplementary Dataset S1) (Fernie *et al*., 2011; Alseekh *et al*., 2021).

### ^13^C-enrichment analysis

The relative abundance of the isotopologues (RIA) (M, M1, M2…Mn) and the fractional ^13^C-enrichment (F^13^C) was calculated as described previously (Lima *et al*., 2018). For instance, in a fragment of three carbons, the sum of the intensity of the isotopologues M, M1, M2 and M3 was set to 100%, and the intensity of each isotopologue is then relative to this sum. In this hypothetical fragment, the F^13^C is calculated according to the following equation: F^13^C = ((M1^*^1)+(M2^*^2)+(M3^*^3))/3. The relative ^13^C-enrichment (R^13^C) was obtained by normalizing the F^13^C by the time 0 of the experiment.

### Metabolic network analysis

Correlation-based metabolic networks were created using metabolite profiling data, in which the nodes correspond to the metabolites and the links to the strength of correlation between a pair of metabolites. The correlation was calculated using debiased sparse partial correlation (DSPC) analysis using the Java-based CorrelationCalculator software (Basu *et al*., 2017). The networks were designed by restricting the strength of the connections to a specific limit of DSPC coefficient (*r*) (−0.5 > *r* > 0.5) using Metscape on CYTOSCAPE v.3.7.2 software (Shannon *et al*., 2003; Karnovsky *et al*., 2012). Network-derived parameters such as clustering coefficient, network heterogeneity, network density and network centralization were obtained as described previously (Assenov *et al*., 2008). We further determined the preferential attachment and the appearance of new hubs, which highlight the nodes that are considered hubs before the start of the experiment (time 0) and maintain its degree of connection and demonstrate nodes that become highly connected after the beginning of the treatments, respectively (Freire *et al*., 2021).

### Statistical analysis

All data are expressed as the mean of four replicates ± standard error (SE). Significant difference in the content or relative ^13^C-enrichment throughout the time was determined by one-way analysis of variance (ANOVA) and Dunnet test (*P* < 0.05), using the time 0 min as control. The difference between dark and light treatments in each time point was determined by Student’s *t* test (*P* < 0.05). These statistical analyses were carried out using SIGMAPLOT12 (Systat Software Inc., San Jose, CA, USA) or Minitab 18 statistical software (State College, PA: Minitab, Inc.). Principal component analysis (PCA) was carried out in square root transformed and auto-scaled (mean-centred and divided by the standard deviation of each variable) data using the Metaboanalyst platform (Chong *et al*., 2018).

## Results

To distinguish dark and light-mediated metabolic changes in guard cells, we performed a ^13^C-HCO_3_ labelling experiment. Guard cells were harvested pre-dawn and subjected to 0, 10, 20 and 60 minutes of continuous darkness or following transfer to white light conditions (400 µmol photons m^-2^ s^-1^). Primary metabolites were identified and used to compare the changes at both metabolite content and ^13^C-enrichment. Whilst changes in the content of metabolites indicate alterations in the pool of them throughout the experiment, the ^13^C-enrichment analysis demonstrates the ^13^C distribution derived from CO_2_ assimilation mediated by PEPc in the dark and by both PEPc and RuBisCO in the light.

### The changes in metabolite content were similar between dark-exposed and illuminated guard cells

Overall, the metabolic changes were similar between dark-exposed and illuminated guard cells. Under both dark and light conditions, the accumulation of fumarate, pyruvate, aspartate and lactate and the degradation of sucrose, glucose, gamma-aminobutyric acid (GABA) and different amino acids was observed (Fig. 1). However, also differential behaviour of specific metabolites was observed. After 60 min of illumination the content of both glutamate (Glu) and urea increased as compared to the time 0 of the experiment and fructose, sorbose and citrate levels decreased (Fig. 1). Comparing dark and light treatments in each time point, decreased content of alanine, citrate, glucose, *myo*-inositol, sorbose and sucrose was observed in illuminated samples. Interestingly, the content of malate is lower at 10 and 60 min and higher at 20 min in the light, as compared to dark-exposed guard cells (Supplementary Fig. S1). This analysis highlights that the dynamic of relative metabolite accumulation/degradation throughout time is slightly altered by light imposition. This idea is further supported by principal component analysis (PCA), in which no clear separation of dark and light treatments was observed (Fig. 2).

**Fig. 1.**
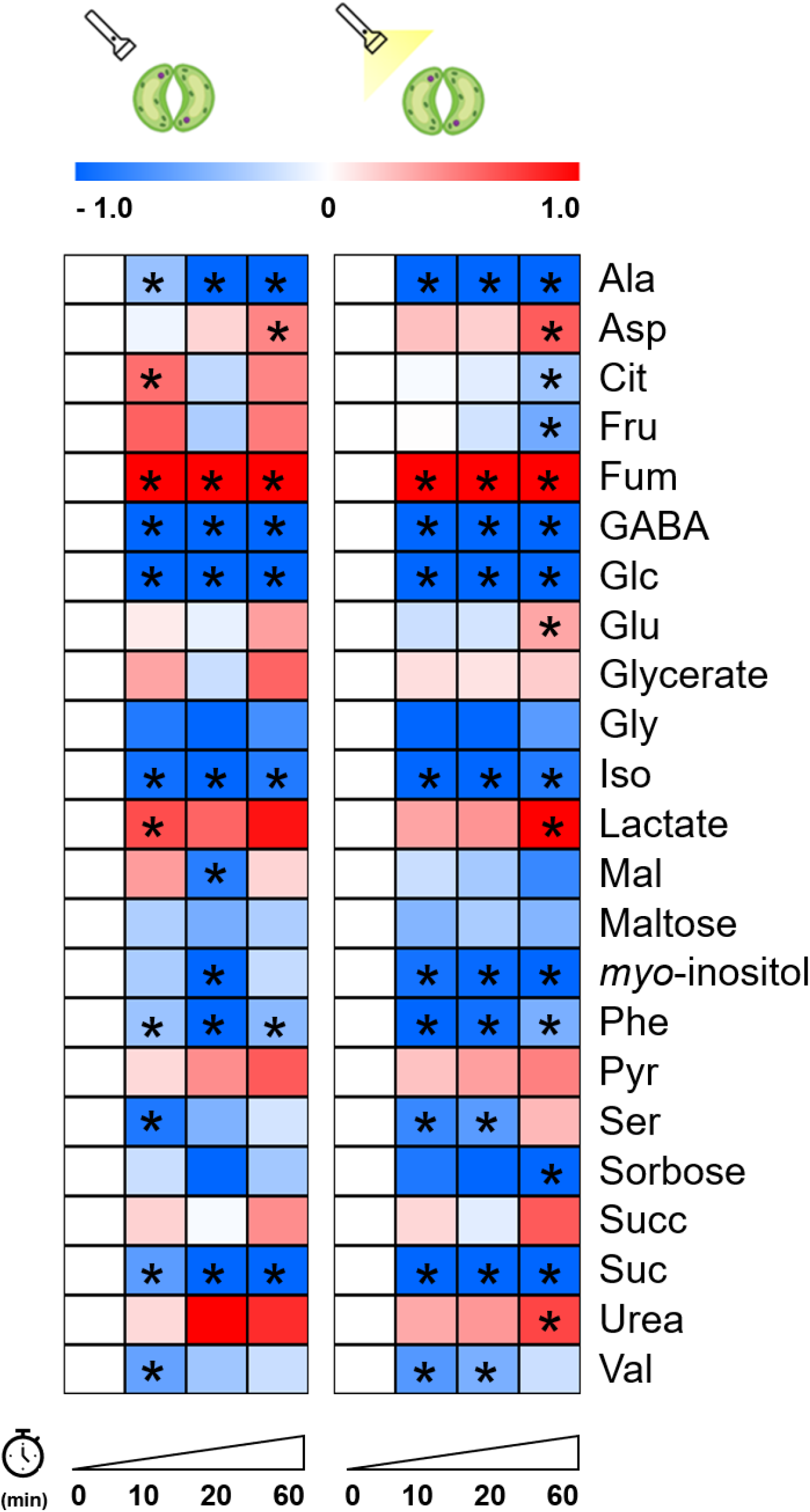
Heat map representation of the metabolite content in guard cells fed with 5 mM of ^13^C-HCO_3_ for 0, 10, 20 and 60 min under dark (left) or light conditions (right). Metabolite content refers to the level of the metabolite normalized by ribitol and the fresh weight used during extraction. The data was further normalized by the values found at the time 0 of the experiment and log_2_-transformed for heat map representation. Asterisks (^*^) indicate significant difference from dark (0 min) by Student’s *t* test (*P* < 0.05).

**Fig. 2.**
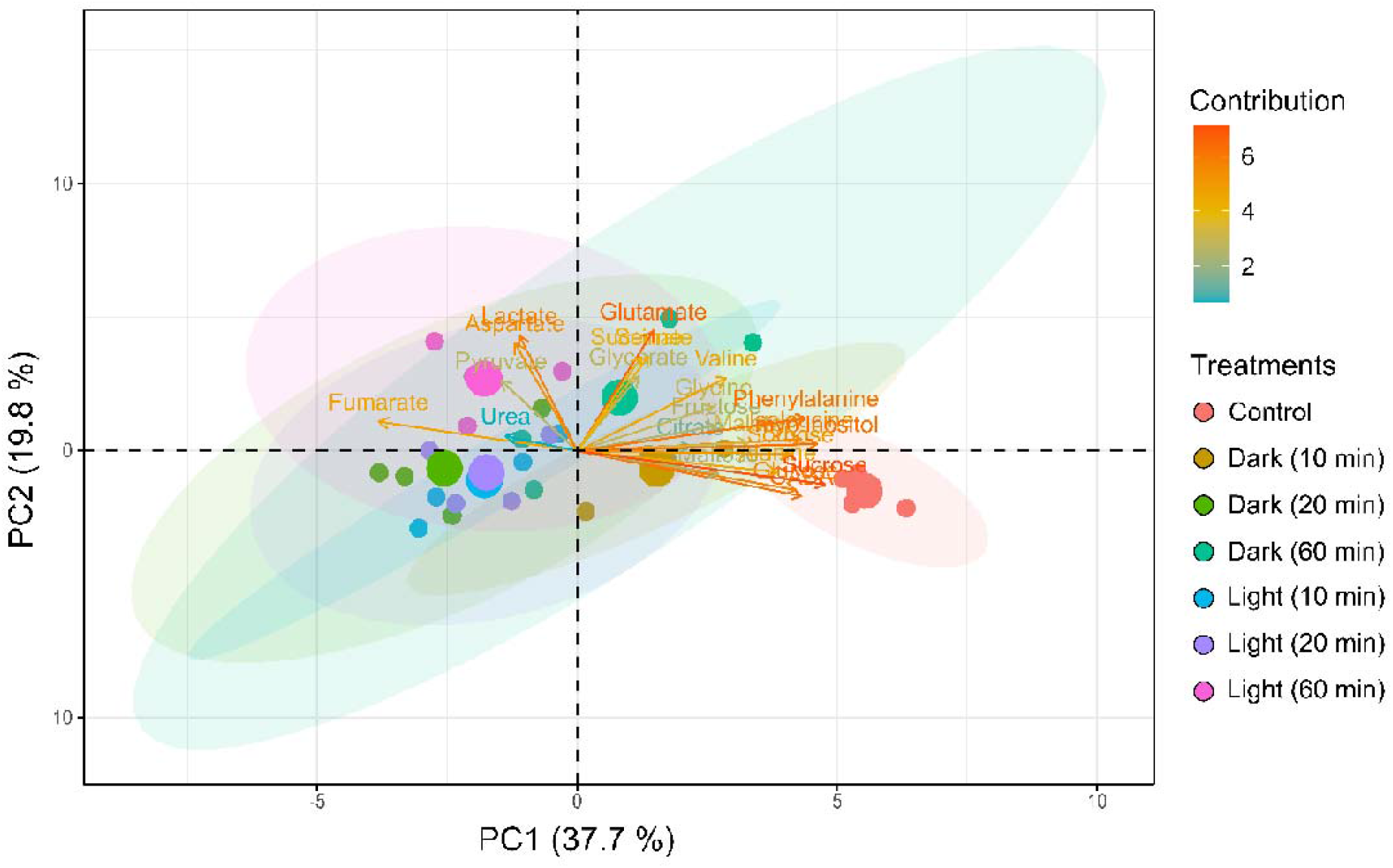
Biplots of the principal component analysis (PCA) of the content of primary metabolites identified from guard cells fed with 5 mM of ^13^C-HCO_3_ for 10 min, 20 min and 60 min under dark or light conditions. The percentage variation explained by the components 1 (PC1) and 2 (PC2) is highlighted between parentheses in each axis. Ellipses surrounding circles of the same colour are the confidence interval of PCA of that treatment. Each colour represents a treatment, as indicated at the right bottom of the figure. The contribution of each metabolite for the distinction between the samples is highlighted according to the colour scale at the upper right of each figure. The contribution of a variable (metabolite) to a given component (PC1 or PC2) is the cos^2^θ of the variable divided by the cos^2^ of the component ×100.

### Illumination triggers re-modelling of metabolic network topology in guard cells

We have recently shown that light-induced stomatal opening involves changes in the density and topology of the guard cell metabolic network (Freire *et al*., 2021). Here, no general pattern of increasing or decreasing network parameters such as clustering coefficient, centralization, density and heterogeneity was observed over time in metabolic networks created using metabolite content data (Supplementary Table S1). However, the number of hub-like nodes, the preferential attachment and the appearance of hub-like nodes differ between dark and light samples. Whilst these parameters reduced to zero after 60 min in the dark, they increased in the light metabolic network over time (Supplementary Table S1), in which malate, fumarate, GABA and glucose appear as important hubs of the metabolic network of illuminated guard cells (Fig. 3). These results suggest that illumination alter the metabolic network structure of guard cells, with particularly strong impact on the pathways associated to the TCA cycle and sugar metabolism.

**Fig. 3.**
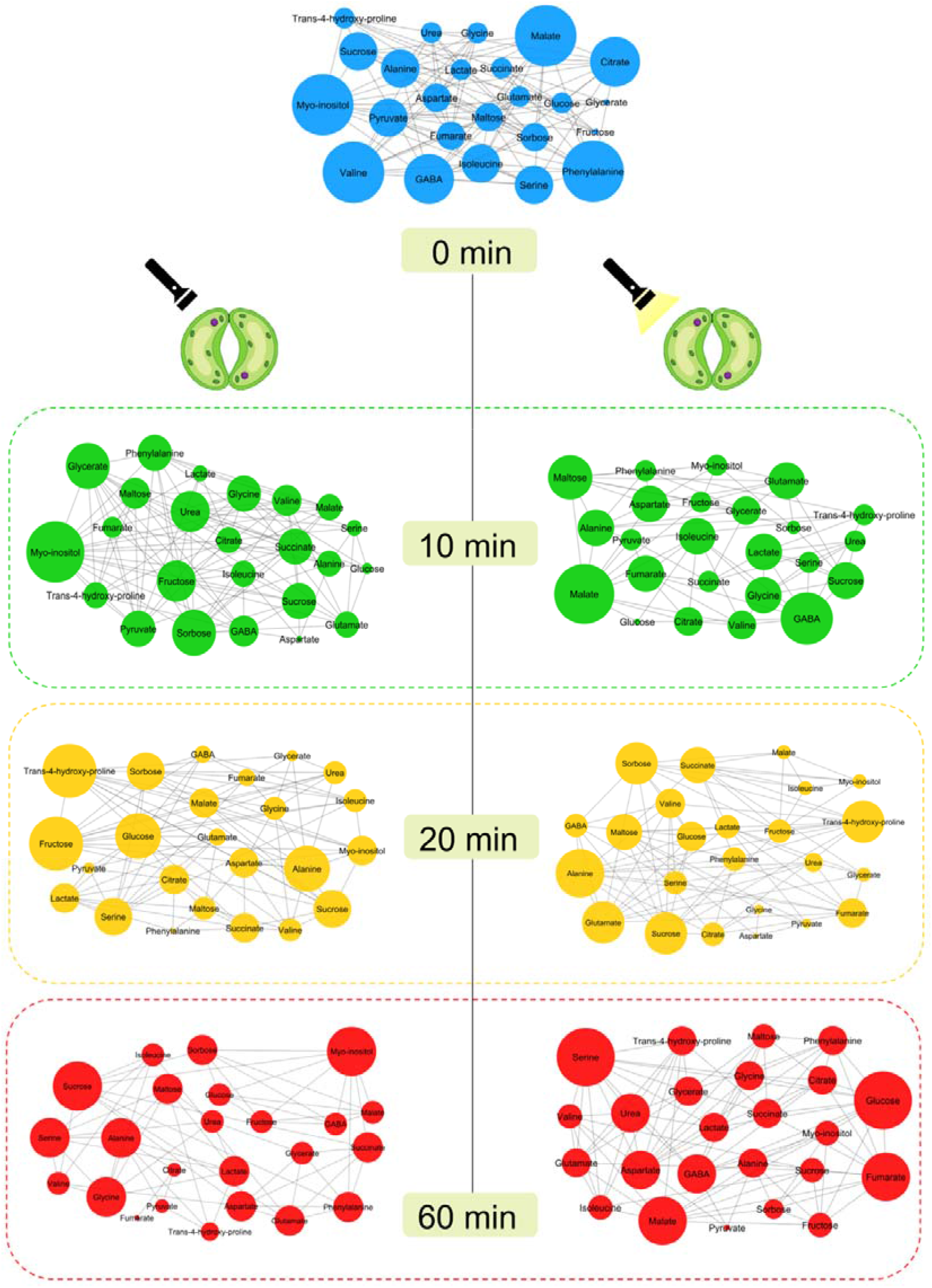
Correlation-based metabolic networks of guard cells fed with 5 mM of ^13^C-HCO_3_ for 0, 10 min, 20 min and 60 min under dark or light conditions. The networks were created using data from relative metabolite content. Nodes are metabolites and links denote debiased sparse partial correlation (−0.5 > *r* > 0.5) between two metabolites. Bigger nodes indicate higher degree of connection, i.e. have more links. These analyses were performed using CorrelationCalculator® and MetScape on Cytoscape®.

### Guard cells have a highly active metabolism in the dark

Similar to the described for metabolite content data, dark-exposed and illuminated guard cells display similar trends in the relative ^13^C-enrichment (R^13^C) data. Alanine, fumarate, glucose, glycine, sucrose and valine were labelled with statistical significance (*P* < 0.05) under both conditions in at least one time point, as compared to the time 0 of the experiment (Fig. 4). However, comparing light and dark treatments in each time point, increased ^13^C-labelling in sucrose, fructose, sorbose, malate, pyruvate, succinate and glutamate was observed in illuminated guard cells, especially at 60 min of labelling (Supplementary Fig. S2). In fact, PCA indicates that dark and light treatments differ mainly at 60 min, as evidenced by the separation by the first component (Fig. 5). Analysis of the contribution of each metabolite for the PCA highlights that certain sugars (sucrose, fructose and glucose), organic acids (citrate and pyruvate), amino acids (Glu, Ile) and *myo*-inositol are those that mostly contributed to the separation observed in PCA of 60 min (Supplementary Fig. S3). These results indicate that several metabolic pathways are activated following PEPc-mediated CO_2_ assimilation in the dark, including gluconeogenesis and the TCA cycle. However, the combination of CO_2_ assimilation catalysed by both PEPc and RuBisCO leads to more dramatic changes in ^13^C-distribution over these and other metabolic pathways in illuminated guard cells.

**Fig. 4.**
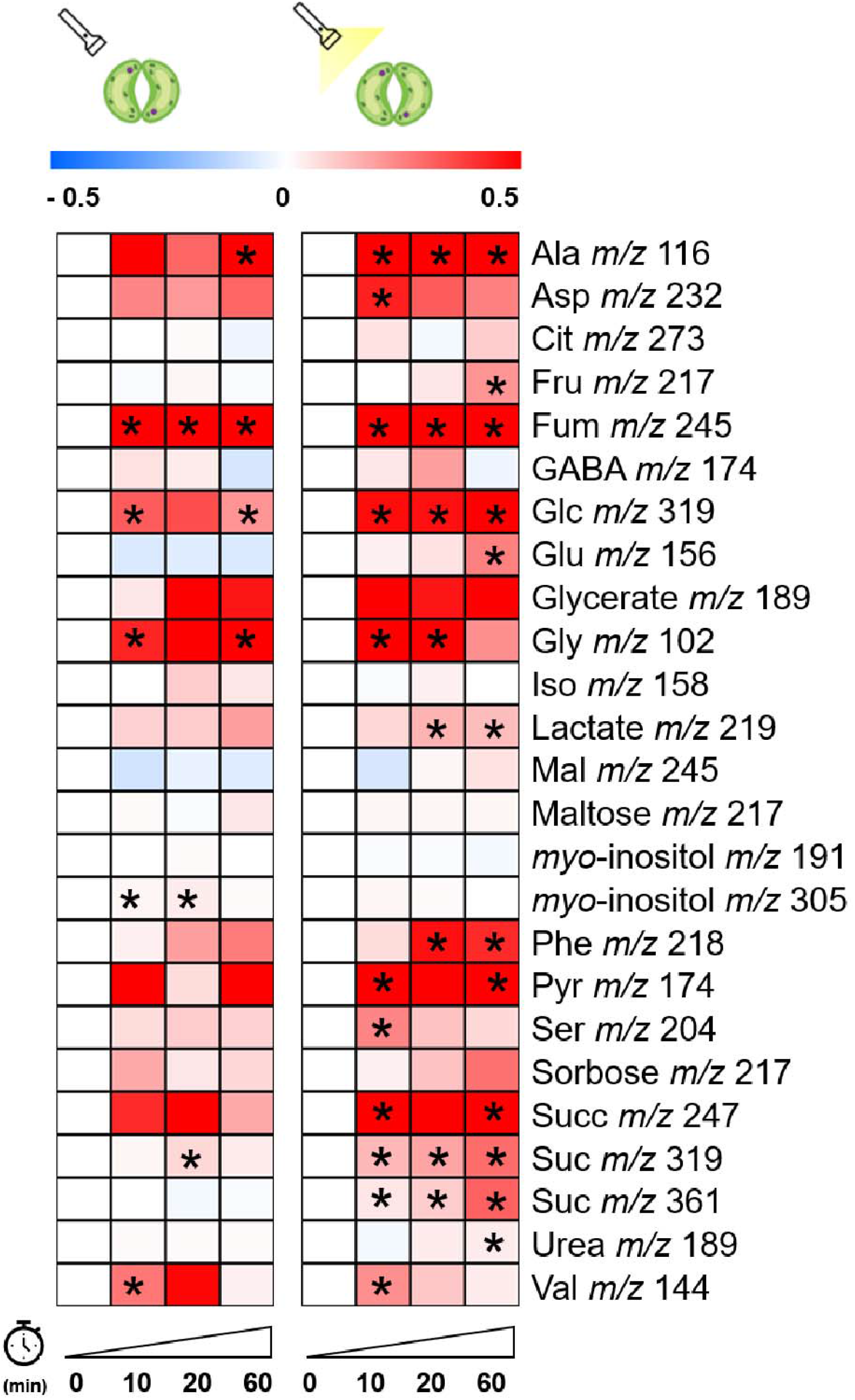
Heat map representation of the relative ^13^C-enrichment (R^13^C) in guard cells fed with 5 mM of ^13^C-HCO_3_ for 0, 10, 20 and 60 min under dark or light conditions. The data was normalized by the values found at the time 0 of the experiment and log_2_-transformed for heat map representation. Asterisks (^*^) indicate significant difference from dark (0 min) by Student’s *t* test (*P* < 0.05).

**Fig. 5.**
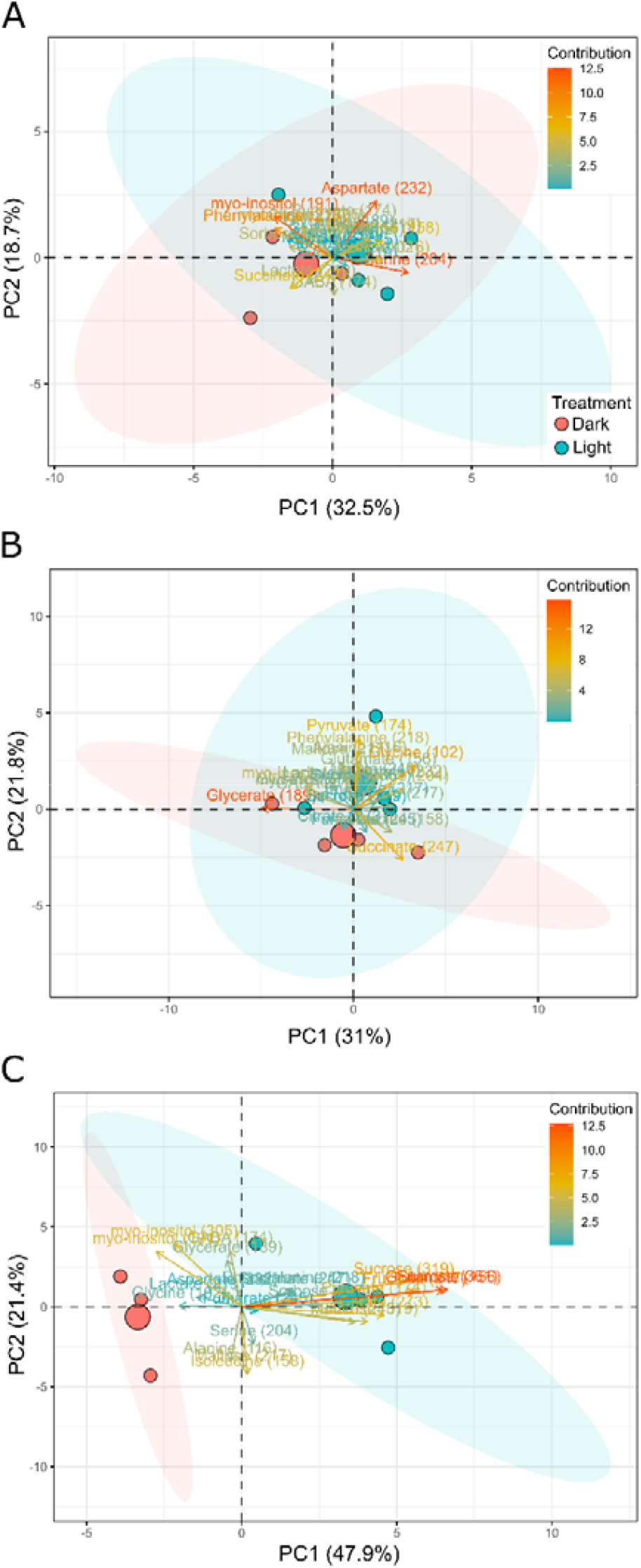
Biplots of the principal component analysis (PCA) of fractional ^13^C-enrichment (F^13^C) in primary metabolites identified from guard cells fed with 5 mM of ^13^C-HCO_3_ for 10 min (A), 20 min (B) and 60 min (C) under dark (red circles) or light (blue circles) conditions. The percentage variation explained by the components 1 (PC1) and 2 (PC2) is highlighted between parentheses in each axis. Red and blue ellipses are the confidence interval of PCA of dark and light samples, respectively. The contribution of each metabolite for dark and light distinction is highlighted according to the colour scale at the upper right of each figure. The contribution of a variable (metabolite) to a given component (PC1 or PC2) is the cos^2^θ of the variable divided by the cos^2^ of the component ×100.

### Fumarate is equally and strongly labelled under both dark and light conditions, but succinate labelling is increased by light exposure

We investigated the R^13^C in the fragments *m/z* 245 of malate, *m/z* 245 of fumarate and *m/z* 247 of succinate, that contain the four carbons (1,2,3,4-C) of these metabolites. Increased R^13^C in fumarate *m/z* 245 under both dark and light conditions was observed (Fig. 4) and no difference in the R^13^C of this fragment was noticed after 60 min of labelling between dark and light conditions (Fig. 6). By contrast, light exposure leads to higher R^13^C in succinate *m/z* 247, when compared to dark-exposed guard cells (Fig. 6). Relative isotopologue analysis (RIA) suggests the incorporation of four ^13^C into succinate under light conditions (Supplementary Fig. S4) (see red circles at the Fig. 6). No significant ^13^C-enrichment in malate *m/z* 245 was observed in either light or dark conditions, when compared to the time 0 of the experiment (Fig. 4). However, the R^13^C-enrichment in malate *m/z* 245 was higher after 60 min in the light, when compared to guard cells kept in the dark (Fig. 6). Intriguingly, the labelling in malate does not match those observed in fumarate and succinate. We have previously shown that only the 4-C of malate was labelled under either dark or light conditions (Lima *et al*., 2021), which contributes to explain the lack of significant labelling in malate *m/z* 245. The labelling in malate could be also interpreted by a dilution in the ^13^C-labelling from previously stored, non-labelled malate, which corroborates the role of malate as a counter-ion of potassium (K^+^) in the vacuole of guard cells (Outlaw and Lowry, 1977; Tallman and Zeiger, 1988; Talbott and Zeiger, 1993).

**Fig. 6.**
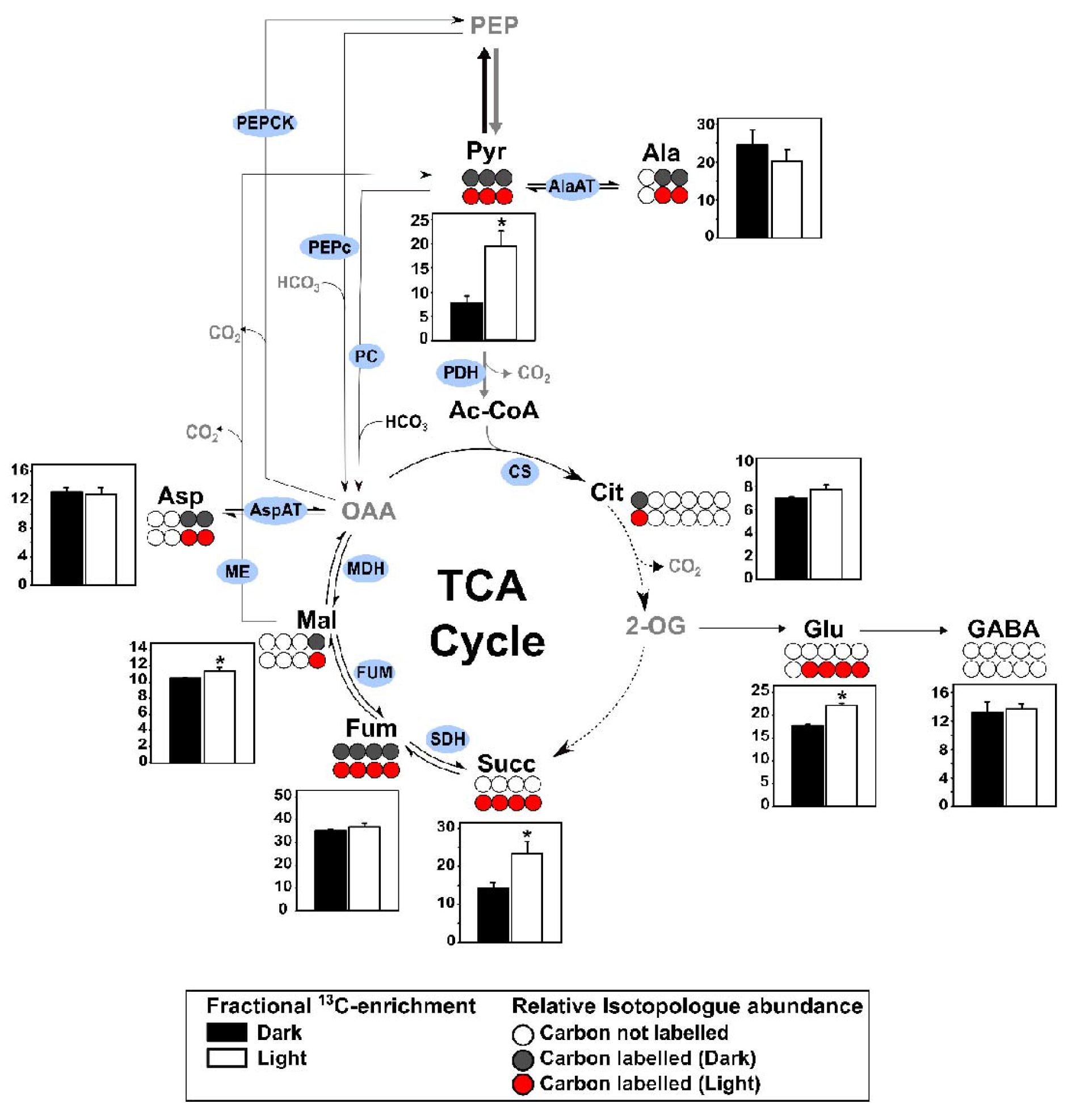
Schematic representation of the fractional ^13^C-enrichment (F^13^C) in metabolites of, or associated to, the tricarboxylic acid (TCA) cycle. Bar graphs demonstrate the F^13^C found after 60 min of 5 mM of ^13^C-HCO_3_ labelling under dark (black bars) or light (white bars) conditions. Spheres are a schematic representation of the relative isotopologue abundance (RIA) data in each metabolite fragment after 60 min of ^13^C-HCO_3_ labelling. RIA indicates the number of labelled carbons incorporated in each metabolite fragment. White spheres represent carbons not labelled, whilst grey and red spheres denote carbons labelled in the dark or light conditions, respectively. Asterisks (^*^) indicate significant difference between dark and light conditions by Student’s *t* test (*P* < 0.05).

### Glutamate synthesis in illuminated guard cells likely depends on PEPc CO_2_ assimilation and previously stored citrate

Illumination leads to increased R^13^C in glutamate *m/z* 156 (Fig. 4). This fragment contains the 2,3,4,5-C of the Glu backbone, which are derived from the TCA cycle (2,3-C) and from pyruvate dehydrogenase (PDH) activity (4,5-C) (Abadie *et al*., 2017). We next investigated the sources of carbon for Glu synthesis in illuminated guard cells by analysing both R^13^C and RIA data of Glu and metabolites of, or associated to, the TCA cycle. Increased R^13^C in pyruvate *m/z* 174 (1,2,3-C) was only observed in the light (Fig. 4). However, increases in the M3 isotopologue (*m/z* 177) of pyruvate was observed at 10 min of labelling in dark-exposed guard cells, indicating the incorporation of three carbons labelled at this time point (Supplementary Fig. S5). After 60 min of labelling, the R^13^C in pyruvate was 2.5-fold higher in the light, when compared to dark-exposed guard cells (Fig. 6). No substantial difference was noticed in the intensity of citrate isotopologues of the *m/z* 273 fragment (1,2,3,4,5-C) in both dark and light samples over time. Although a slight but significant increase in the intensity of *m/z* 274 was noticed (Supplementary Fig. S6), this neither led to an increased R^13^C over time in the dark nor in the light (Fig. 4). Furthermore, no difference in the R^13^C of *m/z* 273 between dark-exposed and illuminated guard cells after 60 min of labelling was observed (Fig. 6). This fragment contains the 1,2,3,4,5-C of the citrate backbone (Okahashi *et al*., 2019). Interestingly, the labelling in citrate does not match those observed in Glu. RIA data of the fragment *m/z* 156 (2,3,4,5-C) indicates that four ^13^C were incorporated into Glu after 60 min in the light, as evidenced by the increases in the isotopologues M1-M4 (*m/z* 157-160) of this fragment (Supplementary Fig. S7). This leads to a higher R^13^C in Glu in the light, when compared to dark-exposed guard cells (Fig. 6). These results suggest that the carbon labelled in citrate is probably derived from PEPc activity (Abadie *et al*., 2017) and that the labelling in citrate is diluted by the incorporation of stored, non-labelled compounds, as previously suggested in leaves (Cheung *et al*., 2014).

Given that Glu could be synthesized by the activity of both Ala and Asp aminotransferases (AlaAT and AspAT) and degraded toward GABA synthesis, we additionally investigated the ^13^C-enrichment in these metabolites. An increased R^13^C in Ala was observed in both conditions, whilst Asp was only labelled in the light, when compared to the time 0 of the experiment (Fig. 4). However, no difference in the fractional ^13^C-enrichment (F^13^C) of these metabolites between dark and light conditions after 60 min of labelling was observed (Fig. 6). No ^13^C-enrichment in GABA *m/z* 174 was observed (Fig. 6). However, assessment of recent data from Arabidopsis guard cells fed with ^13^C-sucrose (Medeiros *et al*., 2018*a*), showed an increased F^13^C in this metabolite during dark-to-light transition (Supplementary Fig. S8). Taken together, these results suggest that whilst both AlaAT and AspAT might contribute to the synthesis of Glu, the reactions catalysed by these enzymes do not fully explain the labelling found in Glu. It seems likely that Glu synthesis in illuminated guard cells mostly depends on carbons from both PEPc-mediated CO_2_ assimilation and previously stored citrate.

### Guard cells have smaller photorespiratory activity than mesophyll cells

Despite the presence of RuBisCO-mediated CO_2_ assimilation in guard cells (Daloso *et al*., 2015*a*), the photorespiratory activity of these cells remains insufficiently understood. To shed light on the relative importance of photosynthesis and photorespiration in guard cells, we next evaluated the ^13^C-enrichment into sugars as metabolites of successful photosynthetic assimilation, and glycine, serine and glycerate as photorespiratory metabolites. Increased R^13^C in sucrose *m/z* 319 (3,4,5,6-C) was observed under both dark and light conditions. However, increased R^13^C in sucrose *m/z* 361 (1,2,3,4,5,6-C) was only observed in the light, when compared to time 0 (Fig. 4). This resulted in a higher R^13^C in both sucrose *m/z* 319 and *m/z* 361 in illuminated guard cells, as compared to dark-exposed guard cells after 60 min of labelling. Similarly, sorbose, fructose and glucose were preferentially labelled in the light (Fig. 7).

**Fig. 7.**
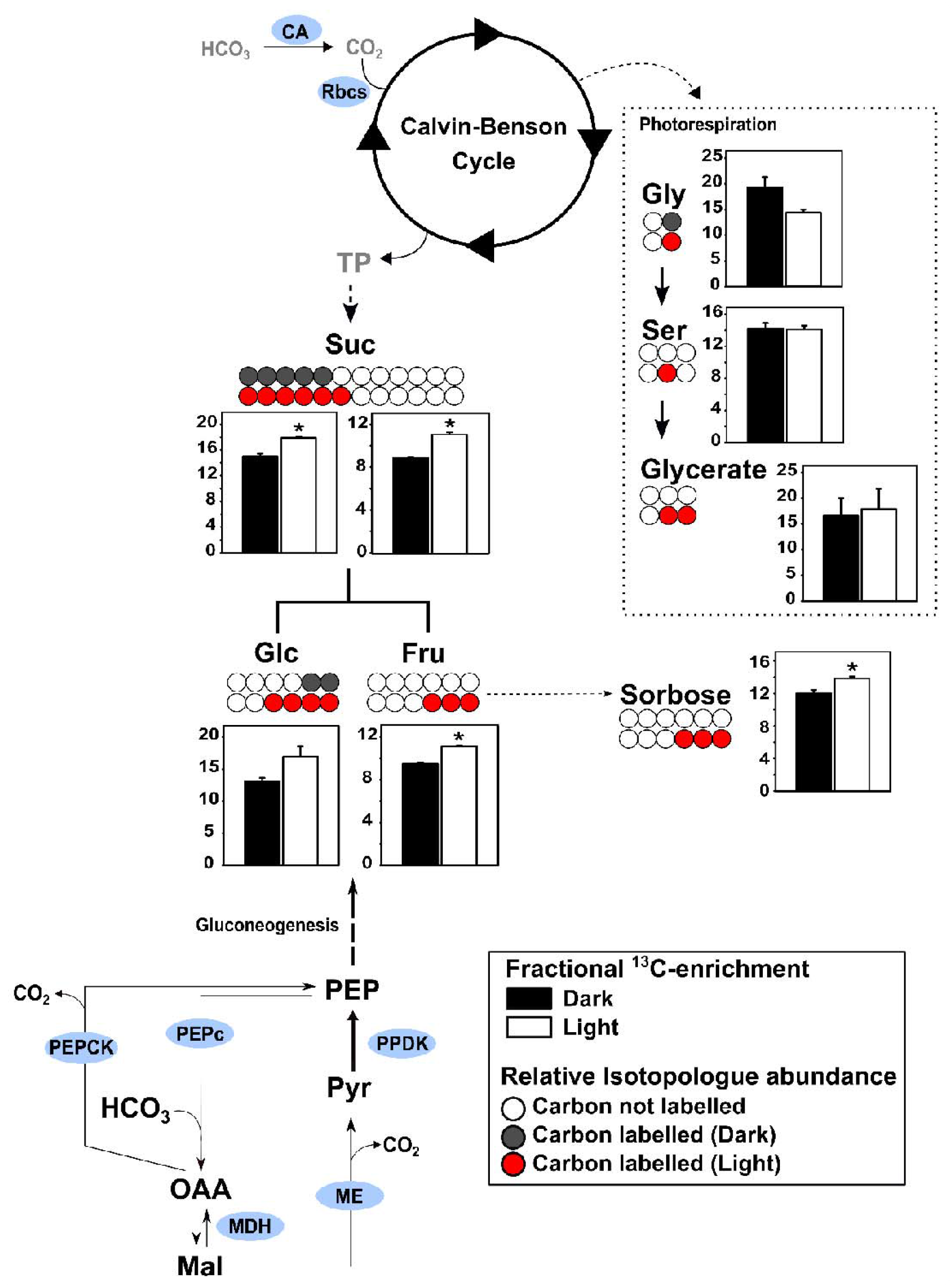
Schematic representation of the fractional ^13^C-enrichment (F^13^C) in sugars and photorespiratory metabolites. Bar graphs demonstrate the R^13^C found after 60 min of ^13^C-HCO_3_ labelling under dark (black bars) or light (white bars) conditions. Spheres are a schematic representation of the relative isotopologue abundance (RIA) data in each metabolite fragment after 60 min of ^13^C-HCO_3_ labelling. RIA indicates the number of labelled carbons incorporated in each metabolite fragment. White spheres represent carbons not labelled, whilst grey and red spheres denote labelled carbons analysed in the dark or light experiments, respectively. Asterisks (^*^) indicate significant difference between dark and light conditions by Student’s *t* test (*P* < 0.05).

Increased R^13^C in glycine *m/z* 102 was observed in both illuminated and dark-exposed guard cells, whilst increased R^13^C in serine *m/z* 204 over time was only observed in the light (Fig. 4). However, no difference in the R^13^C of these metabolites between the treatments after 60 min of labelling was observed. Additionally, glycerate was not labelled in either condition (Fig. 7). We next compared the ^13^C-enrichment in different fragments of both glycine and serine with those obtained in illuminated Arabidopsis rosettes following provision of ^13^CO_2_ (Szecowka *et al*., 2013). The R^13^C in glycine and serine was 2.1 and 4.4-fold higher in whole Arabidopsis rosettes (where mesophyll represents the dominant tissue type) than guard cells (Supplementary Fig. S9), suggesting particularly low photorespiratory activity in guard cells.

## Discussion

### On the complex interplay among the metabolic pathways that regulate the homeostasis of sugars and organic acids in guard cells

Stomata are important regulators of water use efficiency (WUE) in illuminated leaves (Brodribb *et al*., 2019). Furthermore, a recent growing body of evidence suggests that nocturnal stomatal conductance (*g*_sn_) plays an important role in WUE regulation (VialetlChabrand *et al*., 2021; McAusland *et al*., 2021). However, whilst the signalling pathways that regulate stomatal opening in the light have been widely investigated (Inoue and Kinoshita, 2017), the mechanisms that regulate *g*_sn_ remain unknown. Here, we provide insight as to how guard cell metabolism may contribute to the regulation of *g*_sn_. Given that the trends observed in both metabolite level and ^13^C-enrichment are similar between dark-exposed and illuminated guard cells, it could be that similar mechanisms regulate both *g*_sn_ and *g*_s_. This idea is supported by the fact that sucrose breakdown and fumarate synthesis, two mechanisms that support stomatal opening in the light (Daloso *et al*., 2015*a*; Medeiros *et al*., 2016, 2017, 2018*a*; Granot and Kelly, 2019; Flütsch *et al*., 2020*a*), were observed under both dark and light conditions. However, it is important to highlight that the metabolic changes were generally more pronounced in illuminated guard cells. Light exposure may contribute to the activation of several other stomatal regulatory mechanisms in guard cells, given the degradation of starch and lipids in illuminated guard cells (Horrer *et al*., 2016; McLachlan *et al*., 2016; Flütsch *et al*., 2020*a*). Thus, the separation observed in the PCA at 60 min of labelling and the differences in the metabolic networks between dark-exposed and illuminated guard cells could explain the need for more dramatically changes in illuminated guard cells, which would ultimately lead to higher *g*_s_ values, when compared to *g*_sn_. This idea is further supported by the higher degradation rate of sugars observed in the light, which supports findings in which reduced sucrose cleavage capacity of guard cells severely compromises light-induced stomatal opening (Antunes *et al*., 2012; Ni, 2012; Freire *et al*., 2021).

Reverse genetic studies have indicated that alteration in sugar homeostasis in guard cells affects stomatal behaviour (reviewed in Daloso *et al*., 2016b; Flütsch & Santelia, 2021). Additionally, reduced photosynthetic activity in guard cell chloroplasts has been demonstrated to disrupt light-induced stomatal opening (Azoulay-Shemer *et al*., 2015). Although this can been attributed to changes in the cofactor metabolism of ATP and NADPH (Roelfsema *et al*., 2006; Wang *et al*., 2014a), reduced plastidial photosynthetic activity may also compromise sugar homeostasis in guard cells. Indeed, the R^13^C into sugars was higher in the light, evidencing that the RuBisCO-mediated CO_2_ assimilation contribute to sugar synthesis in illuminated guard cells. The ^13^C-labelling incorporation into sugars in the dark suggests that gluconeogenesis is active in guard cells. This corroborates the high ^13^C-enrichment observed in the 3,4-C of glucose under either dark or light conditions (Lima *et al*., 2021), which are proposed to be the glucose carbons preferentially labelled by gluconeogenesis (Leegood and ap Rees, 1978; Beylot *et al*., 1993). These results highlight that gluconeogenesis may be another metabolic pathway that contributes to sugar homeostasis in guard cells, an elusive source of carbon for sugar synthesis in guard cells that has long been debated (Willmer and Dittrich, 1974; Outlaw and Kennedy, 1978; Talbott and Zeiger, 1998; Zeiger *et al*., 2002; Outlaw, 2003; Vavasseur and Raghavendra, 2005; Daloso *et al*., 2016*b*).

Relative isotopologue analysis indicates that three ^13^C were incorporated into pyruvate in both dark-exposed and illuminated guard cells, as evidenced by the significant increases in pyruvate *m/z* 177 after 10 min of exposure to continuous dark or after dark-to-light transition (Supplementary Fig. S5). The labelling in pyruvate in the light might occurs by a combination of ^13^C derived from both RuBisCO and PEPc CO_2_ assimilation, while the labelling in this metabolite in the dark suggests the activity of phospho*enol*pyruvate carboxykinase (PEPCK) and/or malic enzyme (ME), in which labelled OAA and malate would be rapidly converted into PEP and pyruvate, respectively. Additionally, glycolysis and the activity of pyruvate kinase (PK), that converts PEP to pyruvate, could also contribute to pyruvate labelling. Given the labelling observed in sugars in the dark, it seems that the carbon assimilated by PEPc is used to create a substrate cycle between gluconeogenesis and glycolysis, allowing the circulation of carbon between sugars and organic acids without importantly loss of assimilated carbon. Thus, PEPc activity would be important to re-assimilate the CO_2_ lost by several decarboxylation reactions that occurs in chloroplast, mitochondria and cytosol (Sweetlove *et al*., 2013). Indeed, previous modelling results suggest that the flux of CO_2_ from the chloroplast to the cytosol is 17-fold higher in guard cells than mesophyll cells and is largely re-assimilated by PEPc in the cytosol (Robaina-Estévez *et al*., 2017). According to this model, the carbon assimilated by PEPc is transported back to the chloroplast as malate, resulting in a net production of NADPH (Robaina-Estévez *et al*., 2017). These results collectively suggest that PEPc activity is important for both the carbon re-assimilation and the homeostasis of sugars and organic acids in guard cells. The maintenance of a flux of carbon between sugars and organic acids (gluconeogenesis and glycolysis) could be a mechanism to rapidly provide carbons for starch synthesis or for the TCA cycle and associated pathways during stomatal closure and opening conditions, respectively (Outlaw and Manchester, 1979; Medeiros *et al*., 2018*a*).

### Regulation of the TCA cycle and associated metabolic pathways in guard cells

The regulation of plant TCA cycle enzymes is strongly dependent on light quality and quantity (Nunes-Nesi *et al*., 2013). Several lines of evidence point to restricted metabolic fluxes through the TCA cycle in illuminated leaves (Tcherkez *et al*., 2009; Gauthier *et al*., 2010; Daloso *et al*., 2015*b*; Abadie *et al*., 2017; Florez-Sarasa *et al*., 2019), in which different non-cyclic TCA flux modes may contribute to maintain the oxidative phosphorylation system (OxPHOS) (Rocha *et al*., 2010; Sweetlove *et al*., 2010). The GABA shunt has been shown to be an important alternative pathway for the synthesis of succinate, the substrate of the complex II of OxPHOS (Nunes-Nesi *et al*., 2007*b*; Studart-Guimaraes *et al*., 2007). GABA can be synthesized in the cytosol by glutamate decarboxylase or in the mitochondria by GABA transaminases, in which glutamate, 2-oxoglutarate or pyruvate serve as substrates (Bouché and Fromm, 2004). Thus, GABA synthesis is closely associated with the TCA cycle, representing a hub for the C:N metabolic network (Fait *et al*., 2008). Additionally, recent evidence suggests that GABA is an important modulator of stomatal movements, acting as negative regulator of stomatal opening (Xu *et al*., 2021). Our previous data highlighted that the carbon derived from ^13^C-sucrose is incorporated into GABA, but to a lesser extent than into Gln (Medeiros *et al*., 2018*a*). Here, GABA was degraded in a light-independent manner, which may supply OxPHOS with substrate by supporting succinate synthesis in mitochondria. However, no ^13^C-enrichment in GABA was observed. In parallel, increased ^13^C-enrichment in Glu in illuminated guard cells was observed. Therefore, while Glu and Gln seems to be important sinks of the carbons derived from CO_2_ assimilation mediated by both RuBisCO and PEPc, the metabolic flux from Glu/Gln to GABA may be restricted during the dark-to-light transition as a mechanism to allow stomatal opening. This idea is supported by the fact that GABA is a negative regulator of ALTM9 (aluminium-activated malate transporter 9) (Xu *et al*., 2021; Siqueira *et al*., 2021), a key vacuolar anion uptake channel activated during stomatal opening (De Angeli *et al*., 2013; Medeiros *et al*., 2018*b*).

Illumination increased the metabolic fluxes throughout the TCA cycle and associated pathways in guard cells. This idea is supported by the higher F^13^C observed in malate, succinate, pyruvate and Glu in the light, when compared to dark-exposed guard cells (Fig. 6). Furthermore, increased R^13^C in lactate and aspartate was only observed in the light. It is noteworthy that these results were obtained in guard cells with no K^+^ in the medium, given that the presence of this ion strongly increased the ^13^C-enrichment in TCA cycle metabolites, especially in fumarate and malate (Daloso *et al*., 2015*a*). Thus, one would expect that the light and dark metabolic differences may be higher *in vivo*, given that light stimulates the influx of potassium to guard cells (Hills *et al*., 2012; Wang *et al*., 2014*b*,*c*). Surprisingly, the relative content and the R^13^C in fumarate increased substantially under both dark and light conditions. The light-induced ^13^C-enrichment in fumarate resembles previous ^13^C-feeding experiments using ^13^C-HCO_3_ (Daloso *et al*., 2015*a*, 2016*a*; Robaina-Estévez *et al*., 2017). This corroborates the facts that fumarate is the major organic acid accumulated in the light (Pracharoenwattana *et al*., 2010) and that plants with higher fumarate accumulation have higher *g*_s_ (Nunes-Nesi *et al*., 2007*a*; Araújo *et al*., 2011*a*; Medeiros *et al*., 2016, 2017). Furthermore, fumarate emerged as an important hub for the guard cell metabolic network during dark-to-light transition, in agreement with previous observations in guard cells (Freire *et al*., 2021). However, no evidence has hitherto indicated that fumarate is neither the main organic acid accumulated nor can act as osmolyte under dark conditions (Gauthier *et al*., 2010; Araújo *et al*., 2011*b*; Cheung *et al*., 2014; Tan and Cheung, 2020). Thus, whether the accumulation of fumarate in dark-exposed guard cells is a mechanism to sustain *g*_sn_ and/or to store carbon skeletons for the following light period remains unclear. Whilst the characterization of the dynamic of *g*_sn_ in plants with altered fumarate accumulation may be sufficient to understand whether fumarate acts as osmolyte in the dark, testing the second hypothesis will require more sophisticated metabolic experiments to determine the pattern and the subcellular accumulation of fumarate in guard cells during the diel cycle.

Our results suggest that previously stored, non-labelled organic acids are used to support the metabolic requirements of guard cell metabolism in the light, given that the ^13^C-enrichment in citrate and malate is lower than in metabolites of the following steps of the pathway such as fumarate, succinate and Glu. Genome scale metabolic modelling suggests that citrate is the main organic acid accumulated in leaf vacuoles during the night period, which is released and used as substrate for Glu synthesis in the light (Cheung *et al*., 2014). A similar model build specifically for guard cell metabolism predicted that malate accumulates at high rate in the vacuole of guard cells, especially when K^+^ accumulation was restricted by the model (Tan and Cheung, 2020). Taken together, modelling and ^13^C-labelling results indicate the importance of previously stored organic acids to support the TCA cycle and Glu synthesis in illuminated guard cells, resembling the mechanism of TCA cycle regulation observed in leaves (Sweetlove *et al*., 2010; Nunes-Nesi *et al*., 2013; Da Fonseca-Pereira *et al*., 2021).

### On the source of carbons for glutamate synthesis in illuminated guard cells

The connection between the TCA cycle and Glu metabolism is well-established (Araújo *et al*., 2012, 2013; O’Leary and Plaxton, 2020). ^13^C-NMR studies indicate that the synthesis of Glu in illuminated leaves strongly depends on stored compounds (Tcherkez *et al*., 2009; Gauthier *et al*., 2010; Abadie *et al*., 2017), which are presumably accumulated in the previous night period (Cheung *et al*., 2014). This is likely an alternative to overcome the light-inhibition of PDH (Tovar-Méndez *et al*., 2003; Zhang *et al*., 2021), an important source of carbons for the TCA cycle and associated pathways (Reid *et al*., 1977). The degradation of labelled pyruvate by PDH or PC would provide two and three labelled carbons into acetyl-CoA and OAA, respectively. Once synthesized, the acetyl-CoA could be used for citrate synthesis or exported from the mitochondria for fatty acid synthesis (Lonien and Schwender, 2009). However, export of citrate for fatty acid synthesis is unlikely to hold true given that light exposure triggers fatty acid degradation in guard cells (McLachlan *et al*., 2016). It seems likely that fully labelled OAA is synthesized by the activity of both PEPc and PC, which is in turn simultaneously used to the synthesis of succinate through the activity of MDH and fumarase through the C4-branch of the TCA cycle and the synthesis of Glu through the C6(5)-branch of the TCA cycle. In parallel, PDH-derived acetyl-CoA would also contribute to the labelling of Glu through the C6(5)-branch of the TCA cycle.

This labelling pattern throughout the TCA cycle and associated pathways consider the incorporation of non-labelled carbons into malate and citrate. Alternatively, succinate, that is strongly labelled in the light, but not in the dark, could be used as a substrate for Glu synthesis. In this scenario, the TCA cycle would occur in a counterclockwise direction between succinate and 2-OG in illuminated guard cells. Although this possibility has never been confirmed in plants, reverse TCA cycle flux modes have been identified in certain bacteria, being a route to assimilate CO_2_ through the activities of 2-oxoglutarate dehydrogenase (OGDH) and isocitrate dehydrogenase (IDH) in the counterclockwise direction (Evans *et al*., 1966; Buchanan and Arnon, 1990; Mall *et al*., 2018; Steffens *et al*., 2021). Thus, beyond providing carbon skeletons for Glu synthesis, a reverse mode of operation of the TCA cycle would also contribute to increase the CO_2_ assimilated in guard cells. However, this hypothesis is unlikely to occur given the biochemical characteristics of OGDH, that is improbable to occur in the reverse (counterclockwise) direction (Araújo *et al*., 2013), and the fact that this would depend on the activity of 2-OG synthase (E.C 1.2.7.3), that has only been described in bacteria’s (Buchanan and Evans, 1965; Buchanan and Arnon, 1990; Yoon *et al*., 1996, 1997; Dörner and Boll, 2002; Yamamoto *et al*., 2010). Therefore, our results collectively suggest that the carbon derived from PEPc-mediated CO_2_ assimilation is simultaneously used to support gluconeogenesis, the TCA cycle and Glu synthesis and that previously stored citrate and malate may also be an important source of carbons for the TCA cycle and associated pathways.

## Supplementary data

Supplementary data are available at *JXB* online.

Dataset S1. Overview of the metabolite reporting list.

Table S1. Parameters derived from correlation-based metabolic network analysis.

Fig. S1. Heat map representation of the metabolite content in guard cells fed with ^13^C-HCO_3_ for 10, 20 or 60 min under dark or light conditions.

Fig. S2. Heat map representation of the relative ^13^C-enrichment in guard cells fed with ^13^C-HCO_3_ for 10, 20 or 60 min under dark or light conditions.

Fig. S3. Relative contribution the variables for the distinction observed in PCA analysis

Fig. S4. Relative isotopologue abundance of the fragment *m/z* 247 of succinate.

Fig. S5. Relative isotopologue abundance of the fragment *m/z* 174 of pyruvate.

Fig. S6. Relative isotopologue abundance of the fragment *m/z* 273 of citrate.

Fig. S7. Relative isotopologue abundance of the fragment *m/z* 156 of glutamate.

Fig. S8. Fractional ^13^C-enrichment in GABA from guard cells fed with ^13^C-sucrose or with ^13^C-HCO_3_.

Fig. S9. Relative ^13^C-enrichment in glycine and serine in guard cells fed with ^13^C-HCO_3_ and in Arabidopsis rosettes fed with ^13^CO_2_.

## Acknowledgments

This work was made possible through financial support from the National Council for Scientific and Technological Development (CNPq, Grant 428192/2018-1). We also thank the research fellowship granted by CNPq to D.M.D and the scholarships granted by CNPq to F.B.S.F and by the Brazilian Federal Agency for Support and Evaluation of Graduate Education (CAPES-Brazil) to V.F.L. and N.P.P.

## Author contributions

VFL and DMD designed the experiment. VFL, FBSF, NPP and SACS performed the ^13^C-labelling experiment. Mass spectrometry analysis was performed by DBM under the supervision of ARF. Data analysis was performed by VFL, FBSF, SACS, AE, JK and DMD. All authors contributed to write the final manuscript. DMD obtained funding and is responsible for this article.

## Data availability

All data supporting the findings of this study are available within the paper and within its supplementary materials published online.

